# Thermodynamic scaling of canine aging and reversible clock deceleration by a reverse transcriptase inhibitor

**DOI:** 10.64898/2026.02.10.705136

**Authors:** Simon Matrenok, Ekaterina L. Andrianova, Konstantin Avchaciov, Daria I. Fleyshman, Heather J. Huson, John P. Loftus, Joseph J. Wakshlag, Bohan Zhang, Vadim N. Gladyshev, Andrei V. Gudkov, Peter O. Fedichev

## Abstract

Aging was tracked in a cohort of 99 “retired” sled dogs over four years to characterize the latent dynamics of physiological decline. Animals were randomized to receive either placebo or the reverse transcriptase inhibitor lamivudine for ∼30 months. We employed a variational autoregressive model to integrate longitudinal blood parameters and DNA methylation (DNAm) profiles. The model defines Biological Age (BA) as a signature of irreversible damage with Poissonian statistics, a feature conserved across mammalian scales. Lamivudine modulated age-independent latent variables and temporarily decelerated the DNAm clock in females, but these effects were reversible upon treatment discontinuation and did not alter the long-term BA trajectory. Critically, we show that physiological fluctuations are governed by a single systemic factor – an effective or phenotypic temperature representing an emergent (macroscopic) property. We show that while the rate of damage accumulation (the BA slope) is independent of this temperature, actuarial aging parameters (initial mortality and the Gompertz exponent) are strongly temperature dependent. This supports a model where mortality arises from effective activation across a protective free energy barrier that erodes with age. These findings identify phenotypic temperature as an important control variable governing the kinetics of organism-level failure, offering a compelling target for interventions aimed at extending healthspan by “squaring” the survival curve.

## I. INTRODUCTION

Aging of living organisms, particularly mammals, is well described phenomenologically but remains poorly understood at the mechanistic level. On a population scale, aging typically results in an exponential increase in mortality with age—a phenomenon captured by the Gompertz mortality law [1]. On an organismal scale, aging manifests systemically across multiple systems and levels of physiological organization. The diverse manifestations of aging have led to the development of influential frameworks identifying the “hallmarks of aging”: features strongly associated with age and causally linked to the aging process [2, 3]. The number of recognized hallmarks has steadily increased, reflecting the complexity of aging. Moreover, in many systems, these hallmarks are strongly correlated and empirically reversible; interventions targeting certain hallmarks can affect many others. This observation is inherently related to the success of biological clocks—integrated features of the aging process constructed from the statistical averaging of age-related biomarkers [4]. These findings support the prospect of identifying “druggable” targets of aging that could be translated into effective anti-aging therapies. However, the relationship between these microscopic features such as the levels of potentially treatable molecules pathways and the macroscopic outcome of lifespan remains elusive.

Further progress toward clarifying this relationship will require combining in-depth characterization of aging in representative models with testing of targeted interventions, alongside application of advanced statistical tools. Accordingly, in this study, we focused on the aging of domestic dogs maintained under standard conditions and observed during the last third of their lives—a period of progressive systemic physiological decline [5]. Half of the animals were subjected to ∼30 months of treatment with the reverse transcriptase (RT) inhibitor lamivudine, previously shown to inhibit RT activity of LINE-1 (L1) endogenous retrotransposons, whose derepression has been considered a plausible cause of aging-associated inflammation and DNA damage [6]. Among numerous longitudinally recorded health-related traits, we concentrated on the DNA methylation clock, as a representative and objectively quantifiable hallmark of aging [4], and on blood parameters, which in our prior studies have proven to be a reliable and informative source of biological age [7]. By measuring these features in tandem, we show that the key aging signatures are a systemic property that manifests consistently across both molecular (epigenetic) and physiological scales.

Our analysis was built upon our recent proposals suggesting that sufficiently long-living systems can be considered quasistationary systems near thermodynamic equilibrium [8]. Under these circumstances, such systems must allow for statistical descriptions at the level of key thermodynamic variables, such as effective (phenotypic) temperature and “order parameters”—collective emergent variables that characterize the activation of key organism-level system functions. To reverse-engineer the dynamics of the underlying most fundamental variables from noisy observables, we employ a variational autoregressive model—a variant of the structural variational autoencoder—that reveals the signature of aging, represented as a Poissonian statistical random variable—a hallmark we find to be conserved across mammalian species—along with several key health indicators. Using this model, we demonstrate that lamivudine, though previously shown to increase lifespan in a murine model [6], does not affect the biological age variable in dogs. While the drug impacts age-independent features, represented by the latent variables of the model, these effects turned out to be fully reversible and dissipate once the treatment is discontinued.

We show that the fluctuations of the organism’s state variables, represented by the hidden or latent variables of the model, can be statistically described and are predominantly controlled by a single common factor corresponding to an effective phenotypic temperature, a statistical parameter presumably reflecting homeostatic noise [9]. In this framework, the effective temperature serves as a macroscopic state variable that defines the magnitude of fluctuations and, by extension, the organism’s dynamic stability. We find that the rate of aging measured by the slope of biological age as a function of chronological age does not depend on the temperature parameter. However, the actuarial aging parameters, such as the initial mortality rate and the Gompertz exponent, are strongly temperature dependent. This observation supports the hypothesis that mortality results from thermal activation across a protective free energy barrier that progressively declines due to the cumulative effect of accumulated damage, as captured by the biological age variable in the model. We therefore believe that the phenotypic temperature is an actionable variable universally controlling the kinetics of the slowest organism-level changes, such as chronic diseases and risks of death, and hence may be the target of future interventions aimed at increasing healthspan. Our results suggest that while interventions may ‘square’ the survival curve by modulating phenotypic temperature, they do not alter the fundamental structural limit of lifespan of the species.

## II. RESULTS

### A. Survival and DNA methylation clock analyses reveal sex dependence of sled dogs’ aging

The study involved 99 outbred sled dogs of both sexes (54 males and 45 females) collected at an average age of approximately nine years from multiple sled dog kennels. Dogs were maintained in a specialized facility at the Baker Institute of Cornell University (Ithaca, NY) under conditions described in our prior publication [5], including constitutive veterinarian observation, balanced nutrition, a standard vaccination schedule, twice-daily exercise, and periodic analysis of blood parameters, DNA methylation clock (DMC) dynamics, physical conditions, and cognitive functions for four years (Fig. 1a).

**FIG. 1:**
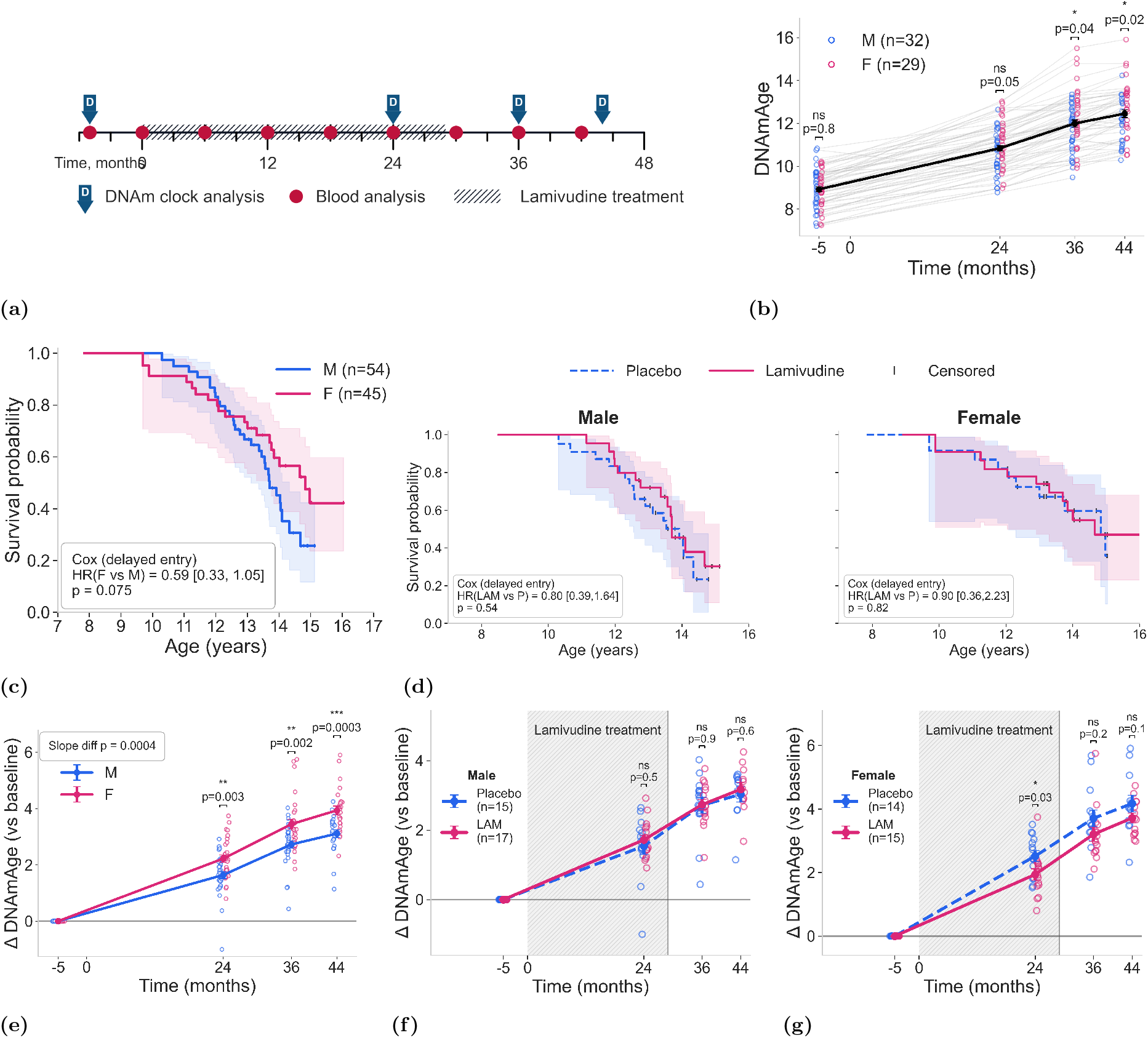
Longitudinal aging and Lamivudine (LAM) intervention in sled dogs. (a) Experimental timeline for 99 dogs over 44 months. Red dots: blood analysis; blue arrows (D): DNAm clock analysis; shaded area: 30-month LAM treatment. (b) Longitudinal DNAmAge progression showing systemic increases across the 44-month window (number of dogs with DNAmAge measured *n*_*M*_ = 32, *n*_*F*_ = 29). (c) Kaplan-Meier survival by sex (*n*_*M*_ = 54, *n*_*F*_ = 45). Females show a non-significant trend toward longevity (*HR* = 0.59, *p* = 0.075). (d) Survival by treatment group. LAM did not significantly alter lifespan in either sex (*p >* 0.5). (e) ∆DNAmAge relative to baseline. Females (red) aged significantly faster than males (blue) in late life (slope difference *p* = 0.0004; month 44 *p* = 0.0003). (f) Male ∆DNAmAge showing no significant LAM effect during (*p* = 0.5) or after (*p*≥ 0.6) treatment. (g) Female ∆DNAmAge showing transient deceleration of epigenetic aging at 24 months (*p* = 0.03), reversing after treatment cessation (months 36–44, *p* ≥ 0.1). Shaded regions in (d, f, g) indicate active LAM administration.

A total of 294 DNA samples isolated from the blood of the studied dogs, which include up to four longitudinal time points per individual spanning four years of observation, were provided to the Mammalian Methylation Clock Consortium [4] for DNA methylation age (DNAmAge) determination. As shown in Fig. 1b, a gradual increase in DNAmAge scores was observed across all longitudinal time points for all dogs.

The experimental design covered the last third of the dogs’ lives, enabling estimation of their longevity. Survival analysis indicates that females live longer than males by *>* 13 months (but given limited sample size the difference was not significant by the standard logrank test, *p* = 0.075, see Fig. 1c), an expected trend that is common for dogs of different breeds as well as for most mammalian species [10, 11].

The dynamics of DNAmAge score increase were steeper in females than in males (Fig. 1e), despite males having a shorter lifespan. Interestingly, the DNAmAge scores were nearly identical at the initial point of observation between males and females, indicating that DMC acceleration in females occurs during the latest period of life. This suggests that DNAmAge score dynamics cannot serve as a biomarker for sex-dependent longevity differences. Similarly, DNAmAge acceleration did not serve as a predictor of mortality rate: Cox models showed no significant association for ∆DNAmAge from the first visit (HR=0.64, p=0.19).

### B. Physiological and epigenetic effects of lamivudine treatment

Treatment of dogs with the reverse transcriptase inhibitor lamivudine was conducted (for the same duration of ∼30 months but starting at different ages ranging from 8.5 to 11.9 years) to test a hypothesis linking aging to the activity of endogenous retrotransposons of the LINE-1 family, previously implicated in aging-related inflammation and the accelerated aging phenotype of Sirt6 knockout mice [6]. Drug dosing was adjusted according to lamivudine pharmacokinetics in dogs to ensure sustained systemic exposure (see Materials and Methods). While no toxicities were detected at the tested dose, supporting the safety of long-term administration in dogs, the drug clearly exerted biological effects on blood markers. To illustrate that, we separated the animals into three groups: non-treated animals, animals receiving treatment, and animals after the treatment (see Fig. S2 with only the features with *p/*15 *<* 0.05 shown).

A spectrum of statistically significant changes in blood biomarkers in lamivudine (LAM) treated dogs collectively point to improved renal function and altered iron metabolism. During treatment, blood urea nitrogen (BUN) and creatinine levels decreased (*p* = 0.003 and *p* = 2.1 × 10^−4^, respectively), suggesting enhanced kid-ney function. Both parameters rebounded after drug withdrawal, with only partial recovery relative to controls (BUN, *p* = 0.15; creatinine, *p* = 0.03), indicating that the beneficial effect was treatment-dependent and reversible. In parallel, iron-related parameters were affected: serum iron and transferrin saturation significantly increased (*p* = 4.6 ×10^−6^ and *p* = 1.9 ×10^−4^), but both returned to baseline after treatment cessation (*p* = 0.27 and *p* = 0.31 vs. controls).

These reversible changes may reflect a reduction in chronic systemic inflammation, which is often associated with altered iron homeostasis and impaired renal clearance [12]. Interestingly, similar shifts in iron and kidney biomarkers have been documented in patients receiving LAM, where they were attributed to suppression of viral replication and consequent reduction in inflammation [13]. Since our study was conducted in uninfected animals, the observed effects may represent a pharmacological action of LAM through inhibition of LINE-1 reverse transcriptase. Additional increase were observed in gamma-glutamyl transferase (GGT, *p* = 4.6 × 10^−4^ during treatment; *p* = 0.051 after cessation), Mean Corpuscular Volume (MCV, *p* = 2.8 × 10^−9^; *p* = 0.14), and Mean Corpuscular Hemoglobin (MCH, *p* = 9.3 ×10^−5^; *p* = 0.11), consistent with previously reported class effects of nucleoside analogs [14]. None of these effects of LAM persisted once the treatment was terminated.

Although lamivudine dosing was sufficient to induce physiological responses, it did not significantly alter lifespan in either sex (Figs. 1d). Nevertheless, we detected the effect of LAM on DNAmAge dynamics: DNAmAge score gain accumulated during ∼30 months of treatment was significantly lower in the lamivudine-treated versus placebo-treated group and reached statistical significance in female dogs, while DNAmAge score dynamics in males were indistinguishable between control and experimental groups (Figs. 1f and 1g). The DNAmAge decelerating trend of lamivudine treatment detected in females disappeared after completion of drug treatment. We concluded that the decelerating effect of LAM on the DNAmAge dynamics in dogs is sex-dependent and, like the effect on individual blood markers, was limited to the period of treatment.

### C. Decoupling Persistent and Reversible Physiological Dynamics

To differentiate between the effects of the drug on aging and the potentially reversible physiological processes, we developed a generative framework designed to disentangle these signals in longitudinal data. We employ a variational structural autoencoder—a generative AI system built to fit longitudinal biomedical data onto the solutions of kinetic equations governing the evolution of a few “hidden” variables. The system consists of three main components: encoder, decoder, and generator models (see Fig. 2). The encoder maps medical inputs (blood tests, age, sex) at any given point in time into a set of probability distributions of latent variables representing the states of underlying systems, which the decoder then maps back to the observables. The generator model uses the latent representations sampled from the encoder to predict their values at a future state. The combined system is trained to minimize the reconstruction loss of the decoder and the Kullback–Leibler (KL) divergence between the probability distributions of the generator prediction and the encoder at every point along the longitudinal health trajectory (see Materials and Methods).

**FIG. 2:**
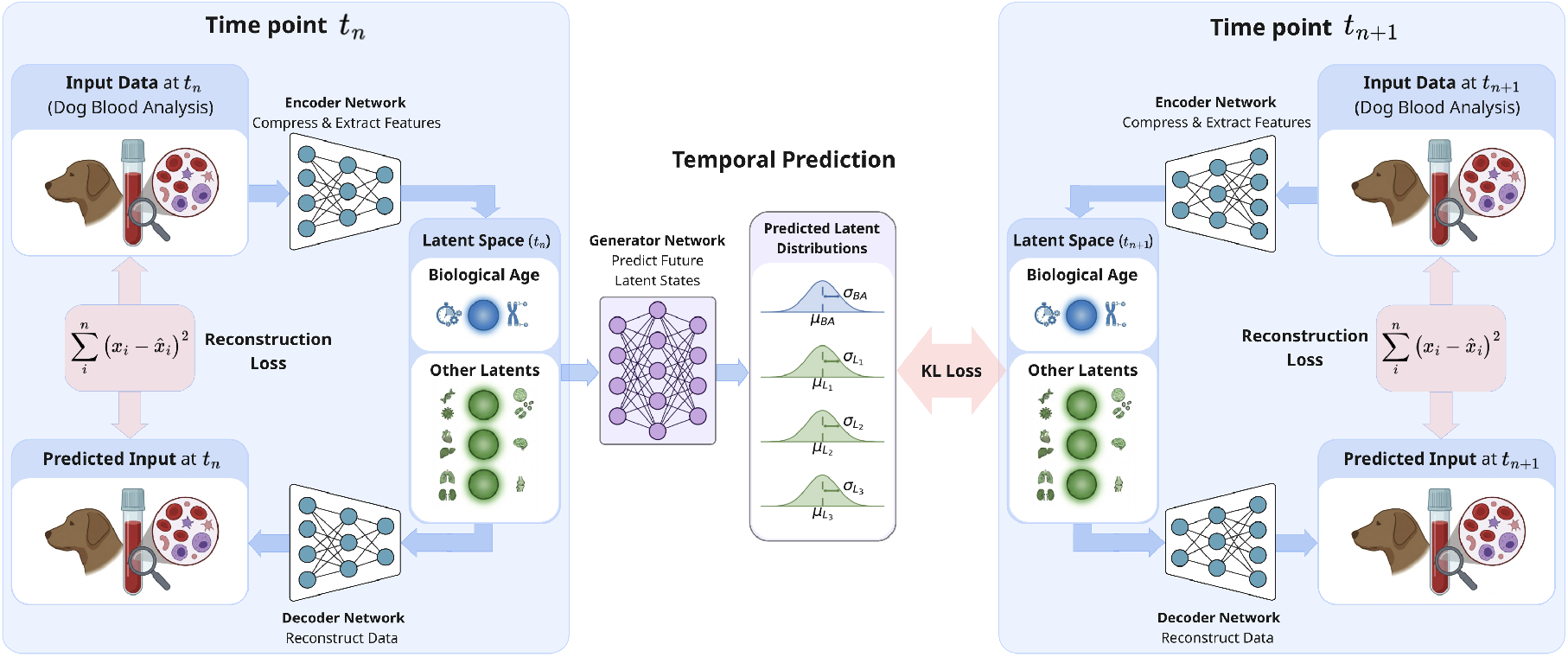
Generative framework for disentangling biological age from transient physiological states. A variational structural autoencoder (VSAE) architecture is applied to longitudinal canine blood data. At time *t*_*n*_, an **Encoder** maps inputs to a partitioned **Latent Space** consisting of a persistent, autocorrelated **Biological Age (BA)** variable and decorrelated physiological latents. A **Decoder** reconstructs the input to minimize error, while a **Generator** predicts the latent transition to *t*_*n*+1_ (**Temporal Prediction**). The model is optimized by minimizing both **reconstruction loss** and the **Kullback–Leibler (KL) Divergence** between predicted and actual latents at *t*_*n*+1_. This dual-objective training isolates deterministic biological progression from stochastic, reversible physiological fluctuations.

In this framework, slow autocorrelated features are assumed to represent slow processes, such as aging, whereas decorrelated features represent stress responses or physiological fluctuations weakly coupled to aging. Accordingly, we introduced a specific latent variable: the biological age (BA) uniquely identified by its long autocorrelation property, whereas all other latent variables are presumed to be decorrelated along the life history of the animals.

Once trained, we first examined the properties of the biological age derived from the model. Fig. 3a illustrates the averaged trajectory of BA segmented into three phases: pre-treatment (including the control group, shown in blue), active Lamivudine (LAM) treatment (magenta), and post-treatment (green). Our results show no discernible effect of LAM treatment on the overall BA trajectory, indicating that the drug does not significantly impact the persistent latent signal associated with aging. Consistent with the hallmarks of a stochastic damage accumulation process, we observed that both the mean and the variance of the BA increase linearly with chronological age (Fig. 3b). Furthermore, BA acceleration—defined as the excess of BA over the mean in an age-matched cohort—is significantly associated with an elevated risk of death (Cox proportional hazard model *HR* = 1.48, 95% *CI* = 1.17–1.87, *p* = 0.002).

**FIG. 3:**
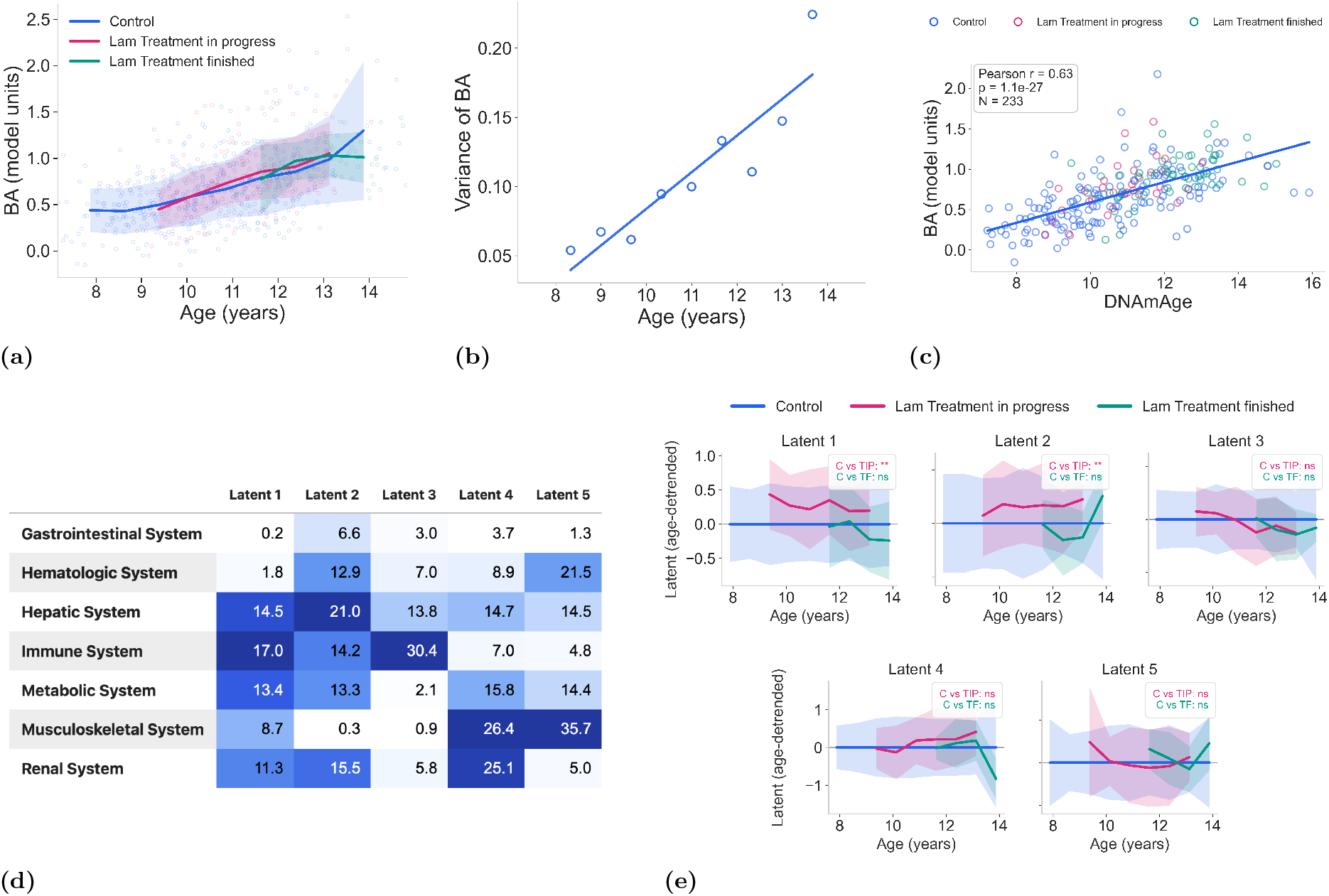
Latent space characterization: persistent aging vs. reversible physiological states. (a–b) **Properties of Biological Age (BA):** (a) Average BA trajectories during control (blue), active LAM treatment (pink), and post-treatment (green) phases. The lack of significant deviation suggests LAM does not alter the fundamental rate of damage accumulation; (b) Variance of BA increases linearly with age; (c) **External Validation:** Correlation between model-derived BA and DNA methylation age (DNAmAge) (*r* = 0.63, *p* = 1.1 × 10^−27^), confirming the latent variable tracks biological aging. (d) **Systemic Associations:** Heatmap of associations between model latents and clinical systems. Values represent −log_10_(*p*-value) of the correlation between latent scores and blood chemistry markers, grouped by functional system. Higher values (darker blue) indicate stronger statistical evidence for the association. (e) **Pharmacological Response:** Age-detrended trajectories for reversible latents. Significant shifts occur in Latent variables 1 and 2 during active treatment (** *p <* 0.01 vs. control) but return to baseline upon cessation (ns). This demonstrates the model’s capacity to isolate transient pharmacological effects from the persistent aging process.

Moreover, the BA derived from the model correlates strongly with DNA methylation clocks (Pearson *r* = 0.6, *p* = 10^−22^), a correlation that remains significant even after de-trending for common dependence on chronological age (Pearson *r* = 0.47, *p* = 4 ×10^−9^), see Figs. 3c and S5b, respectively. These results suggest that while lamivudine may influence specific biomarkers and epigenetic scores, it does not alter the fundamental, persistent latent variable associated with the organism’s irreversible aging process.

### D. Biological meaning of the latent variables and the effects of treatment

The variational autoregressive model identified five latent variables (labeled 1 through 5) that capture distinct physiological processes and individual health indicators (see Figs. 3d and S3, where we show the linear associations of the latent variables and the observable features—individual measurements).

Latent Variable 1 highlights a combined axis of protein balance, immune cell activity, and electrolyte regulation. Its strongest associations include total protein, globulin, potassium, monocyte count, eosinophil count, phosphate, and the sodium-to-potassium ratio. Taken to-gether, these features suggest that these latent variable captures activation of the immune system, with a predominant contribution from innate immune responses. The increase in monocyte and eosinophil counts points directly to activation of innate immune cell populations, while the shifts in protein fractions (total protein and globulin) are consistent with enhanced production of acute-phase and immunoglobulin proteins during inflammation. In parallel, alterations in electrolytes and phosphate, as well as the sodium-to-potassium ratio, likely reflect systemic adjustments secondary to immune activation and metabolic stress. Thus, Latent Variable 1 can be interpreted as a composite signature of innate immune activation accompanied by metabolic and electrolyte remodeling.

Latent variable 2 is associated with iron transport, red blood cell characteristics, and kidney function. Notable features include globulin, albumin-to-globulin ratio, FE saturation, creatinine, blood urea nitrogen (BUN), and red blood cell indices. The combination of iron-related measures (FE saturation) and erythrocyte indices (MCV, MCH) points to a close relationship with red blood cell function and iron utilization. At the same time, the inclusion of creatinine and BUN suggests that renal function contributes to this axis, consistent with the kidney’s central role in erythropoiesis and iron metabolism through erythropoietin production and iron recycling. Protein parameters (globulin, albumin-to-globulin ratio) may further reflect the systemic balance between nutritional status and chronic inflammation, both of which influence erythrocyte homeostasis. Altogether, Latent Variable 2 can be interpreted as a composite marker of red blood cell physiology and iron metabolism.

Latent Variable 3 aligns with liver enzyme activity and leukocyte involvement. Key contributors include the hepatic markers ALT, AST, AP, and GGT—enzymes that typically reflect hepatocellular integrity or biliary function [15]—together with elevated signals from WBC counts and monocytes, indicative of immune modulation. A strong loading on bicarbonate may further suggest links to systemic acid–base balance.

Latent Variable 5 is predominantly driven by red blood cell parameters and markers of tissue turnover. It features especially large loadings for hematocrit, hemoglobin, lactate dehydrogenase (LDH), and CK, as well as anion gap. Clinically, this latent variable is notable because it is the only one associated with an elevated risk of death (Cox proportional hazard model *𝓏* = 2.13, *CI* = 0.02–0.38, *p* = 0.03), implying that RBC dysregulation or systemic energy metabolism captured here may predict poorer survival outcomes.

Lamivudine administration had a transient impact on Latent Variables 1 and 2 only (see Fig. 3e). These shifts were evident only during the course of treatment and reverted to baseline afterward, mirroring a similar reversible pattern observed in the DNAm Clock measures.

### E. Thermodynamics of biological noise and Gompertz law

The longitudinal nature of the dataset allows us to investigate the properties of biological fluctuations and their relationship to all-cause mortality. Specifically, we used repeated measurements along each animal’s life trajectory to examine the statistical properties of fluctuations in the model’s latent variables in relation to remaining lifespan and the pattern of mortality acceleration with age.

We calculated the variance for each latent variable (excluding biological age) across each animal’s trajectory. Matrix factorization via Singular Value Decomposition (SVD) revealed that a single factor explains ≈80% of this biological noise. This confirms the existence of a single, pathway-independent scaling factor controlling the power of physiologically relevant noise within the cohort, having the meaning of the “effective temperature”, a key finding from the cross-species analysis in [9]. To support this argument, we immediately observed that the variance of DNAmAge residuals (calculated after accounting for age and treatment) also significantly correlates with the effective temperature derived from our latent variables (Fig. 4a).

**FIG. 4:**
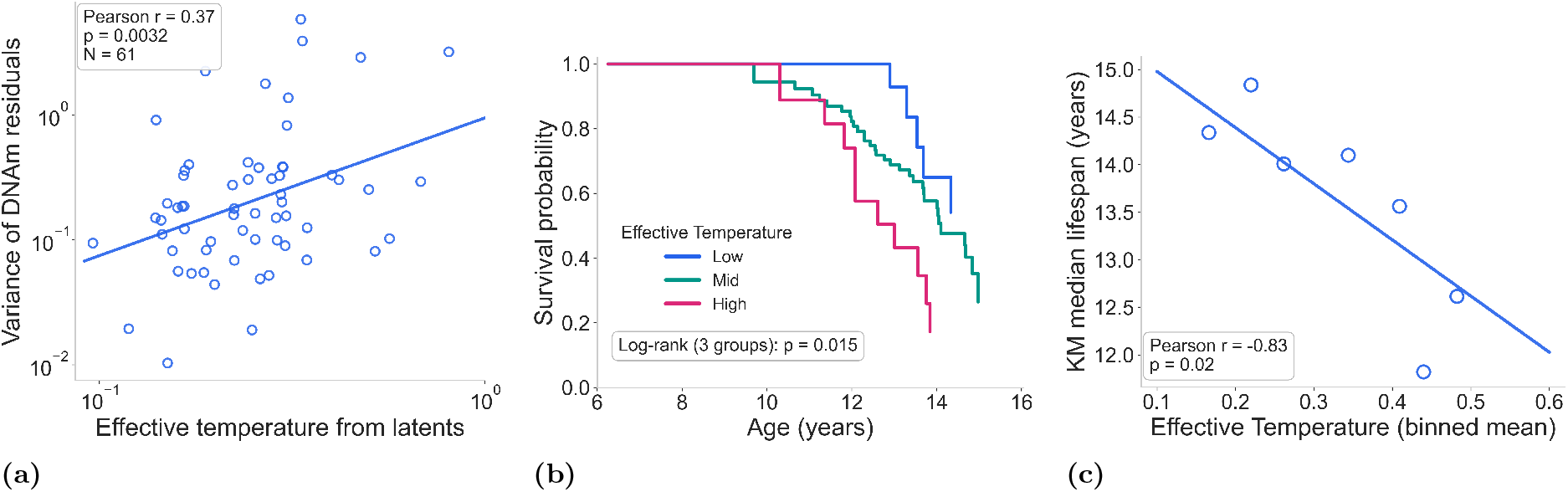
Multiscale manifestation of phenotypic temperature and its impact on mortality. **(a)** Correlation between the effective temperature derived from physiological latents and the variance of DNA methylation age (DNAmAge) residuals (*r* = 0.37, *p* = 0.0032, *N* = 61). This significant association indicates that thermodynamic noise manifests consistently across physiological and epigenetic scales; **(b)** Kaplan-Meier survival curves for canine cohorts stratified into terciles based on their effective phenotypic temperature derived from the latent state model. The Low Temperature cohort (blue) exhibits the highest survival stability, maintaining a rectangular survival profile until the structural biological limit, whereas the High Temperature cohort (red) shows significantly earlier onset of mortality; **(c)** Correlation between the mean phenotypic temperature lifespan in cohorts binned by the effective temperature.

To evaluate the biological implications of the novel state variable, the average variance of all latent variables at each time point was utilized as a proxy for the effective phenotypic temperature (*T*) of the animals. We first observed that the damage accumulation rate, measured by the slope of biological age (BA) as a function of chronological age, showed no dependence on phenotypic temperature (Fig. S4a).

Survival analysis was performed on the study population (*N* = 99) stratified by temperature strata, representing the bottom, middle, and top bins of the effective phenotypic temperature distribution. Kaplan-Meier analysis demonstrated a statistically significant separation in mortality risk across these cohorts (Log-rank *p* = 0.01, Fig. 4b). The “coldest” cohort (*T*_*low*_) exhibited a rectangularized survival profile with mortality concentrated near a structural ceiling of approximately 170 months. In contrast, the “hottest” cohort (*T*_*high*_) showed a progressive acceleration of death events, significantly reducing the median survival age compared to the lower-temperature groups.

Average lifespan decreased linearly as a function of the phenotypic temperature index. Weighted linear regression across eight temperature-stratified cohorts yielded a slope of approximately −30 months per unit of effective temperature (Fig. 4c). The difference in lifespan between the extreme cohorts (representing the lowest and highest temperature octiles) was approximately 3 years. The regression intercept of approximately 15.5 years represents the theoretical structural limit for this population (see Discussion).

The dynamics of mortality were further characterized by examining the mortality hazard across temperature-stratified cohorts (Fig. S4b). Utilizing 15-month binning, a distinct separation in the baseline and trajectory of death rates was observed between the phenotypically “cold” and “hot” populations. Gompertzian modeling indicated that dogs in the high-temperature group established a higher mortality risk profile beginning as early as 100 months of age. Conversely, the actuarial aging rate (*α*) was steeper in the lower-temperature group compared to the high-temperature group. The mortality risk curves for both groups converged at approximately 15 years, consistent with the structural limit identified in Fig. 4c.

## III. DISCUSSION

Our study, which monitored “retired” sled dogs [5] during the last third of their lifespan, was initiated with two major goals: (i) to determine whether inhibition of endogenous reverse transcriptase activity exerts an antiaging effect in canines, and (ii) to generate new insight into large-animal aging through deep longitudinal profiling of nearly 100 dogs maintained under standardized husbandry conditions. Here we present findings addressing both objectives.

The rationale for testing reverse transcriptase inhibitors as anti-aging agents is grounded in multiple observations implicating endogenous retrotransposons in aging pathogenesis – mediated by reverse transcriptase encoded by the LINE-1 family [16]. These elements, which collectively occupy nearly half of mammalian genomes, are epigenetically silenced in healthy cells but become derepressed in senescent cells [16]. Epigenetic derepression of LINE-1 due to Sirt6 gene knockout drives chronic inflammation and persistent DNA damage, phenocopying accelerated aging in mice [6]. Remarkably, germline retrotransposition is uniquely suppressed in the exceptionally long-lived naked mole rat, suggesting a role of retrotransposon control in extreme longevity [17, 18]. Pharmacological inhibition of LINE-1 reverse transcriptase by nucleoside RT inhibitors such as lamivudine mitigates LINE-1–driven pathology in Sirt6-deficient mice and blocks LINE-1-mediated tumor adaptation to chemotherapy [6]. Moreover, the extraordinary phenotypic diversity of dog breeds, largely driven by mobile element insertions, reflects high retrotransposon activity in canines [19]. Collectively, these findings support the hypothesis that RT inhibition may attenuate aging processes in dogs.

In our study, half of the dog cohort received lamivudine at a dose and treatment schedule selected to achieve sustained systemic exposure. Multiple physiological indicators demonstrate lamivudine’s effects in vivo consistent with what was observed in anti-HIV treated patients. All observed drug-related reactions were fully reversible upon treatment cessation and were not associated with any toxicities. Over ∼30 months of continuous administration, lamivudine did not significantly extend lifespan in either sex; however, it produced a clear decelerating effect on the DNA methylation clock in females. Similar to many other geroprotective interventions, this putative anti-aging effect appeared sex-specific [10, 11]. This effect was evident during treatment and disappeared after withdrawal, along with physiological effects of lamivudine exposure.

The epigenetic signal observed with lamivudine—specifically, deceleration of DNAmAge scores in females—supports continued investigation of reverse transcriptase (RT) inhibition as a potential gerotherapeutic strategy. The absence of a stronger or more sustained physiological effect may reflect pharmacokinetic (PK) limitations, such as insufficiently high or stable plasma lamivudine concentrations to achieve robust and continuous inhibition of LINE-1 RT activity. Consistent with this interpretation, lamivudine PK in dogs appeared less favorable than in humans, prompting us to adjust dosing levels and administration schedules (twice-daily dosing) in an effort to maintain adequate drug exposure (see Materials and Methods). Future studies should focus on further optimization of dosing regimens and on evaluation of more potent RT inhibitors with improved PK properties. Importantly, the favorable safety profile observed with long-term lamivudine administration in dogs further supports the feasibility of RT inhibition as a therapeutic approach for aging-related interventions.

To differentiate reversible drug- and stress-induced fluctuations from progressive, age-dependent factors within our longitudinal dataset, we developed a generative framework designed to disentangle these distinct signals. Building upon and extending our previous work [7], we employed a Variational Structural Autoen-coder (VSAE) – a specialized variant of the structural variational autoencoder architecture [20]. This deep neural network is optimized to solve two problems simultaneously: first, it compresses the high-dimensional biological state into a minimal set of independent latent variables; second, it identifies their underlying trajectories by fitting the latents to kinetic equations of motion governing the state space.

To capture the underlying aging dynamics, we introduced a specific latent variable, Biological Age (BA), characterized by high longitudinal autocorrelation. This property uniquely separates BA from other latent variables, which appear decorrelated over subsequent measurements and likely represent transient physiological fluctuations. We observed that BA increases progressively with age, alongside a concordant linear rise in its variance, which provides strong evidence for its nature as a Poissonian random variable. This Poissonian signature is not unique to canine physiology; rather, it appears to be a universal feature of mammalian aging. Similar characteristics have been identified in DNA methylation (DNAm) profiles in mice [21] and humans [22], as well as in a broad cross-species analysis spanning 348 evolutionarily distant mammals [9]. In all cases, this feature is fundamentally tied to the accumulation of entropic damage [22, 23].

The strong correlation between our model-derived BA and the DNAmAge score reinforces this connection. Crucially, the persistence of this relationship after regressing out the shared dependence on chronological age demonstrates that the model captures a meaningful biological signal beyond simple time-tracking. This cross-scale correlation—linking macroscopic physiological parameters to molecular epigenetic states—supports the interpretation of BA as a thermodynamic and emergent state variable. In this framework, BA represents a systemic (or macroscopic) property that manifests consistently across multiple biological subsystems.

The remaining latent variables of the model represent unique aspects of biological health, reflecting different systems or processes involved in physiology, ranging from immune response and oxygen transport to electrolyte balance and organ function. By applying this model, we confirmed that while Lamivudine impacts age-independent physiological features – specifically Latents 1 and 2 – it does not alter the trajectory of the BA latent. This insight is inherently lost when biological complexity is condensed into a single DNAmAge clock readout. Instead, our framework provides a more nuanced view by successfully disentangling functional health systems from the irreversible process of aging.

The model assumes that latent variables other than BA are decorrelated across subsequent measurements. Such short autocorrelation times imply that these features represent systems in stable equilibrium governed by relaxational dynamics, which inherently predicts the transient character of responses to external perturbations [7]. The transient nature of the observed drug effects—which dissipated upon treatment cessation—suggests that the impact on maximum lifespan may be negligible if the intervention is terminated before the end of life. This pattern of dynamic stability is reminiscent of the reversible impact of smoking on biological age observed in humans [24].

Furthermore, the lack of response of the stochastic aging signature (BA) to pharmacological intervention is consistent with findings in murine models, where neither caloric restriction nor heterochronic parabiosis successfully altered the stochastic damage signature [21]. While Lamivudine appears to modulate specific immune-metabolic or renal pathways, its failure to intercept the BA trajectory in dogs highlights a critical distinction between entropic damage and other dynamic, potentially reversible physiological state variables. Although our current data does not permit a direct mapping of BA to configurational entropy directly, this observed resilience to intervention supports the interpretation of BA as a quantitative indicator of a thermodynamically irreversible aging process.

The longitudinal character of our dataset allows us to demonstrate that physiological fluctuations represented by the model’s latent variables are predominantly governed by a single systemic factor. This provides a compelling instance of emergent thermodynamic properties in an aging organism, a phenomenon we previously identified in a cross-species analysis of DNA methylation spanning 348 mammalian species [9]. Such behavior is characteristic of complex systems operating near a state of dynamic stability and serves as a manifestation of the fluctuation-dissipation theorem [25] which establishes a fundamental proportionality between the magnitude of fluctuations (biological noise) and the recovery rates of individual physiological pathways (dissipation; see e.g., [24]). In this context, the system-level, process-independent proportionality coefficient functions as an “effective temperature”.

This effective temperature represents a universal property of biological noise at time scales relevant to lifespan determination and is distinct from kinetic body temperature [9]. Like Biological Age (BA), it is an emergent macroscopic property; when estimated from clinical biochemical factors, as in this study, it correlates with the variance of the DNAmAge score residuals (after regressing out the effects of age, sex, and treatment). Consequently, it defines the common factor behind the power of the noise across specific physiological subsystems.

As a macroscopic property, the effective temperature is alongside BA a key determinant of longevity. Its role can be understood through a basic stability analysis (see the summary in [8]). The short autocorrelation times of all latent variables other than BA suggest that young animals exist in a state of dynamic stability. In this regime, physiological fluctuations are stochastic and correspond to relaxation dynamics in the presence of ran-dom forces, representing the internal and external stress factors jointly described by the effective temperature.

The basin of attraction has a finite radius, and the probability of an activation transition representing the loss of dynamic stability and subsequent death defines the mortality rate at an age *t, M* (*t*). This risk is modeled as: *M* (*t*) = *κ* exp(−*U/T*) where *U* is the activation barrier, *T* is the effective temperature, and *κ* is the “attempt frequency.” The attempt frequency represents the time scale characterizing the activation and relaxation rate of the most relevant process with the smallest activation barrier (more details can be found in the Kramer’s theory section in [25]).

Because aging is a slow process occurring on time scales that vastly exceed those of individual physiological processes, the risk of an activation transition depends on macroscopic quantities: the effective temperature (*T*) and the biological age (BA). This dependence is mediated through the regulatory free energy, *U* (BA) ≈*U*_0_ −*U* ^′^BA, where *U* ^′^ = *dU/d*BA. Given that BA increases linearly with age (BA = *R*_*D*_*t*), the damage accumulation rate *R*_*D*_ is independent of the effective temperature, as demonstrated here by comparing aging rates across animals of the same species (Fig. S4a) and previously across 348 mammalian species [9].

Accordingly, the risk of death follows the Gompertz law, *M* (*t*) = *M*_0_ exp(*αt*), where the initial mortality rate is *M*_0_ = *κ* exp(−*U*_0_*/T*) and the actuarial aging rate (the Gompertz exponent) is: *α* = *U* ^′^*R*_*D*_*/T*. The aging rate is thus proportional to the damage accumulation rate and inversely proportional to the effective temperature. Consequently, the average lifespan (*t*_ls_) is a linear function of the effective temperature:

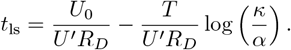

Since aging is slow relative to individual processes (*κ ≫ α*), the logarithmic term is large (and hence weakly temperature dependent) and always positive. In this framework, the effective temperature acts as a “fluctuation tax,” reducing the average lifespan relative to the structural limit corresponding to the maximum lifespan (the first term and regression intercept). This exactly matches the linear dependence observed in our data (Fig. 4c).

The effective temperature therefore controls the discrepancy between the average and the maximum lifespan—the point at which the basin of attraction and dynamic stability (resilience) are lost. This explains why the effective temperature is responsible for the “squaring” of the survival curve (Fig. 4b). Understanding the biology governing *T* could, in principle, allow us to bridge the gap between the current (∼80 years) and maximum (∼120 years [24]) lifespan in our own species—a hefty goal that makes effective temperature a plausible target for pharmacological, behavioral or dietary interventions aimed at extending healthspan.

A temperature-like variable in relation to the Gompertz law was first proposed by Strehler and Mildvan in [26]. This work, alongside our recent analysis [9], represents the first successful identification of this parameter from non-demographic, clinical or molecular level biomedical data. Beyond providing support for the theoretical thermodynamic model of aging, the increasing availability of rich genetic and molecular datasets should enable the identification of specific targets that control the effective temperature.

We therefore propose that phenotypic temperature is a potentially actionable variable that controls the kinetics of the slowest organism-level changes, such as chronic disease progression and mortality risk. As such, it is a primary target for future interventions aimed at increasing healthspan by squaring the survival curve. The maximum lifespan, however, remains temperature-independent and awaits the discovery of additional regulatory control variables and may require radical engineering solutions.

## Appendix S1: Materials and methods

### 1. Animals

The study cohort comprised retired sled dogs recruited from multiple kennels across North America. This population was selected for its high genetic diversity and consistent prior life history as high-performance athletes, minimizing confounding effects associated with breed-specific pathologies. The cohort consisted of clinically healthy dogs with no severe underlying conditions requiring chronic systemic therapy at the time of enrollment. All subjects were relocated to the Vaika research facility at the Cornell University College of Veterinary Medicine to ensure environmental and nutritional consistency. The dogs were maintained under uniform, climate-controlled conditions with continuous professional veterinary oversight and were fed a standardized diet (Annamaet Extra 26). This centralized housing and husbandry model was implemented to reduce environmental and lifestyle variability typically associated with community-dwelling pet populations, thereby providing a stable physiological baseline for longitudinal analyses. All experimental protocols were approved by the Institutional Animal Care and Use Committee (IACUC) of Cornell University’s College of Veterinary Medicine. Details of the colony establishment, maintenance, and care have been described previously [27].

#### 2. Lamivudine

The nucleoside reverse transcriptase inhibitor lamivudine is an oral drug approved for the treatment of patients with HIV [28]. To determine appropriate dosing and administration frequency, we analyzed the pharmacokinetics (PK) of lamivudine in six randomly selected sled dogs from our colony. Plasma lamivudine concentrations were quantified using a validated highperformance liquid chromatography–tandem mass spectrometry (HPLC-MS/MS) assay. Blood samples were collected into Na_2_-EDTA tubes and stored at −80 ^◦^C until analysis.

Sample preparation involved protein precipitation with acetonitrile: 200 µL of plasma was mixed with 600 µL of acetonitrile containing an internal standard. Following centrifugation at 13,500 rpm for 10 minutes at 4.0 ^◦^C, the supernatant was evaporated to dryness under nitrogen at 37 ^◦^C. The residue was reconstituted in 1,000 µL of water containing 0.1% formic acid.

Chromatographic separation was performed using an HPLC system coupled to tandem mass spectrometric detection. A 10.0 µL aliquot of the reconstituted sample was injected for analysis. The assay was calibrated over a concentration range of 0.1–10 µg/mL, with sensitivity and specificity meeting standard bioanalytical validation criteria for the detection of lamivudine in canine plasma. PK analysis indicated a lamivudine plasma half-life of approximately 2–2.5 hours in dogs, compared with 5–7 hours in humans. Based on these findings, lamivudine was administered twice daily (rather than once daily as typically used in humans) at a dose of 10 mg/kg to achieve more sustained systemic exposure.

#### 3. Trial design

The data presented in this investigation were derived from a large-scale, prospective, randomized, open-label, placebo-controlled longitudinal trial conducted in a cohort of retired sled dogs. Detailed descriptions of the parent study recruitment procedures and baseline assessments have been reported previously [27]. While the parent study encompassed a broad range of physiological and behavioral endpoints, the present analysis focused specifically on survival outcomes, longitudinal hematological parameters, and changes in DNA methylation patterns (see Fig. S1).

A total of 104 dogs aged 8–11 years were screened, of which 99 were enrolled and randomized by age and sex. The experimental group (n = 50; 23 females, 27 males) and the placebo group (n = 49; 21 females, 28 males) were housed in a dedicated facility under standardized conditions. The treatment phase lasted 30 months (April 2019 to October 2021), with total monitoring extending to 50 months (concluding in June 2023).

Dogs in the experimental group received oral lamivudine at a dose of 10 mg/kg twice daily for 30 months (April 2019 to October 2021). The control group received a placebo identical in size and weight. Individual doses were periodically adjusted based on semiannual body weight measurements to maintain target exposure throughout the treatment phase. Following completion of the 30-month treatment period, drug administration was discontinued and animals entered a 20-month follow-up observation phase. Survival status was monitored continuously throughout the study period to assess all-cause mortality.

#### 4. Blood collection

The current analysis included cellular and biochemical parameters assessed in blood samples collected at nine time points: screening, baseline, and months 6, 12, 18, 24, 30, 36, and 42. Baseline blood collection was performed no earlier than one month after animal relocation to the facility to ensure stabilization of housing, dietary, and environmental conditions across the study cohort. Venous blood samples were obtained by jugular venipuncture following an overnight fast to minimize postprandial metabolic variability. Samples were collected into EDTA tubes for hematological analysis and DNA extraction, and into serum separator tubes for biochemical profiling. All samples were processed on the day of collection at the Cornell University College of Veterinary Medicine clinical laboratories, with aliquots stored at −80 ^◦^C for subsequent longitudinal DNA methylation analyses.

#### 5. DNA methylation clock analysis

DNA was isolated from the whole blood of 61 dogs selected from healthy animals for whom four longitudinally collected samples were available with similar representation of sexes within placebo- and lamivudine-treated groups. DNA methylation profiling was conducted at four timepoints, including baseline, and months 24, 36, and 42. Altogether, 294 DNA samples were analyzed.

DNA methylation–based age estimation was performed using epigenetic clocks developed by the Mammalian DNA Methylation Consortium [29]. Genomic DNA was isolated from dog samples using standard column-based purification methods, followed by quality and quantity assessment by spectrophotometry and fluorometry. Only samples with high molecular weight DNA and A260/280 ratios between 1.8 and 2.0 were used for downstream analysis.

Genome-wide DNA methylation profiling was conducted using the HorvathMammal methylation array (Illumina), which interrogates conserved CpG sites across mammalian species. Bisulfite conversion of genomic DNA was performed according to the manufacturer’s protocol, followed by array hybridization and scanning at a certified service provider affiliated with the DNA Methylation Clock Consortium. Raw intensity data (IDAT files) were processed using the consortium’s standardized bioinformatics pipeline. Briefly, data were background-corrected, normalized, and filtered for quality control using established preprocessing procedures. DNA methylation levels were expressed as *β* values ranging from 0 (unmethylated) to 1 (fully methylated).

Epigenetic age estimates were calculated using previously published dog-specific and pan-mammalian DNA methylation clock algorithms, as appropriate. DNAm-Age acceleration or deceleration was defined as the residual from linear regression of DNAmAge on chronological age, thereby accounting for age-dependent baseline methylation changes. Sex-specific analyses were performed when indicated. All epigenetic clock calculations and statistical analyses were performed using R software with scripts and coefficients provided by the Mammalian DNA Methylation Consortium. Investigators performing methylation analyses were blinded to experimental group assignments where applicable.

#### 6. Statistical analysis

Statistical modeling was tailored to the longitudinal nature of the extracted data. Linear mixed-effects models were utilized to analyze changes in blood parameters over time, accounting for repeated measures within individual subjects. Survival distributions were estimated using the Kaplan-Meier method, and differences between groups were assessed using the log-rank test. The impact of the intervention on longevity was further quantified using a Cox proportional hazards model.

#### 7. Data preparation

The dataset comprised longitudinal records from 99 dogs, each with up to nine sequential blood parameter measurements. The interval between consecutive measurements was approximately six months. Each measurement included 43 features, which were normalized using a power transformation.

In total, we had 104 dogs enrolled in the study. However, 2 dogs were excluded due to missing date of birth information, and 3 additional dogs died prior to randomization, leaving 99 dogs in our final dataset. Of these, DNA methylation (DNAmAGE) data was available for 61 dogs, which is why some analyses are based on this subset.

#### 8. Model architecture and training

The Variational Structural Autoencoder (VSAE) consists of three main components: the Encoder, Decoder, and Generator.

##### Encoder

The Encoder receives blood parameter measurements along with normalized age and sex as inputs. It maps these inputs into a latent space composed of two parts: a biological age (BA) estimator and a set of physiological latent variables. As in a standard Variational Autoencoder (VAE), the Encoder predicts distributions (means and variances for a Gaussian distribution) for each latent variable, from which values are sampled during training.

##### Decoder

The Decoder takes the sampled BA and physiological latents and reconstructs the blood parameter measurements. This encoding and decoding process is applied independently at each available time point in the trajectory for every dog.

##### Generator

To capture temporal dependencies, the Generator models the evolution of latents across time points. It takes the current BA estimate and physiological latents as inputs and predicts the expected distribution of these variables at the next time point.

A key architectural feature is the distinct treatment of temporal autocorrelation for BA and physiological latents:

- The BA estimator is explicitly designed to be autocorrelated across adjacent time points—that is, the BA estimate at one time point is expected to be similar to (and therefore predictable from) the BA estimate at the previous time point. Intuitively, this reflects the idea that biological age should evolve smoothly over time rather than change abruptly. The Generator enforces this by predicting the next BA as a Gaussian distribution whose mean and variance depend on the current BA value.
- In contrast, physiological latents are designed to exhibit little or no autocorrelation across time points. These latents are intended to capture relatively high-frequency or short-term biological variations, such as transient physiological fluctuations, measurement noise, or unmodeled environmental factors that can change rapidly and unpredictably. Because the interval between blood measurements is relatively long, we assume that any specific shortterm state captured by these latents at one measurement is unlikely to persist to the next, resulting in low or zero autocorrelation. The Generator therefore predicts the next physiological latents using a Gaussian distribution whose parameters do not depend on their previous values, but may still depend on the current BA. This approach enables the model to account for age-dependent changes in physiology while not artificially linking transient effects across time points.

##### Training Procedure

The Encoder, Decoder, and Generator are trained jointly. The loss function combines two terms:

- Reconstruction loss between the observed blood measurements and those reconstructed by the Decoder.
- Kullback-Leibler (KL) divergence between the distribution of latents inferred by the Encoder at each time point, and the expected latent distributions predicted by the Generator for that time point based on the latents from the *previous* time point. This encourages the latents inferred by the Encoder to be consistent with the temporal dynamics learned by the Generator.

For the first time point in each trajectory, where no prediction is available from the Generator, the KL divergence is computed with respect to a state-independent prior that may still depend on age.

##### Effective temperature computation

The purpose of the effective temperature is to quantify, with a single scalar value per dog, how strongly the physiological latent variables fluctuate over time. Conceptually, this quantity is analogous to temperature in physics: higher temperature corresponds to larger random fluctuations of microscopic states. In our setting, a higher effective temperature reflects larger temporal variability of the physiological latent representation.

##### Latent fluctuation profiles

For each dog, we first quantify how much each physiological latent varies across the available time points. Specifically, for dog *i* and physiological latent dimension *k*, we compute the variance across time:

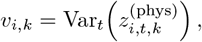

where 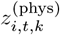 denotes the inferred value of the *k*-th physiological latent at time point *t*.

This procedure yields a matrix

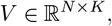

where *N* is the number of dogs and *K* is the number of physiological latents. Each row of *V* represents a dog-specific “fluctuation profile,” describing how strongly each latent dimension varies over time for that dog.

##### Dominant mode of variability across dogs

To understand whether these fluctuation profiles differ in a complex, multidimensional way or mainly by a single underlying factor, we perform singular value decomposition (SVD) of the matrix *V* . SVD identifies orthogonal modes that explain decreasing amounts of variance across dogs.

We observe that the first singular value is much larger than the remaining ones:

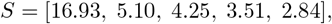

and that the first mode alone explains approximately

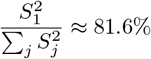

of the total variance in *V* .

This indicates that, to a very good approximation, differences between dogs can be described by a single factor that scales the magnitude of fluctuations across *all* physiological latents simultaneously. In other words, dogs primarily differ not in *which* latents fluctuate, but in *how strongly* they fluctuate overall.

##### Effective temperature definition

The coordinate of each dog along this first SVD mode provides a natural one-dimensional summary of its latent fluctuations. We interpret this scalar as the dog’s *effective temperature*: a single parameter capturing the overall intensity of physiological variability across time.

##### Practical estimator used in this study

We further find that the effective temperature obtained from the first SVD mode is highly correlated with a simpler quantity: the average variance of physiological latents for each dog,

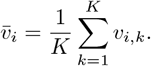

Because of this strong correspondence, we use 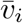 as the effective temperature estimate in our analyses. This estimator is straightforward to compute and retains the same biological interpretation as the SVD-based temperature.

## Appendix S2: Supplementary Figures

**FIG. S1:**
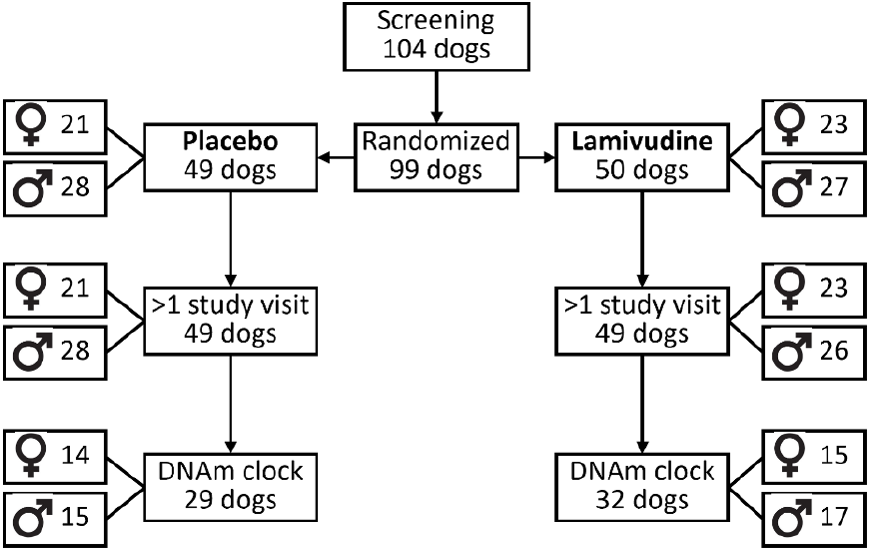
Scheme of distribution of dogs among the study cohorts.

**FIG. S2:**
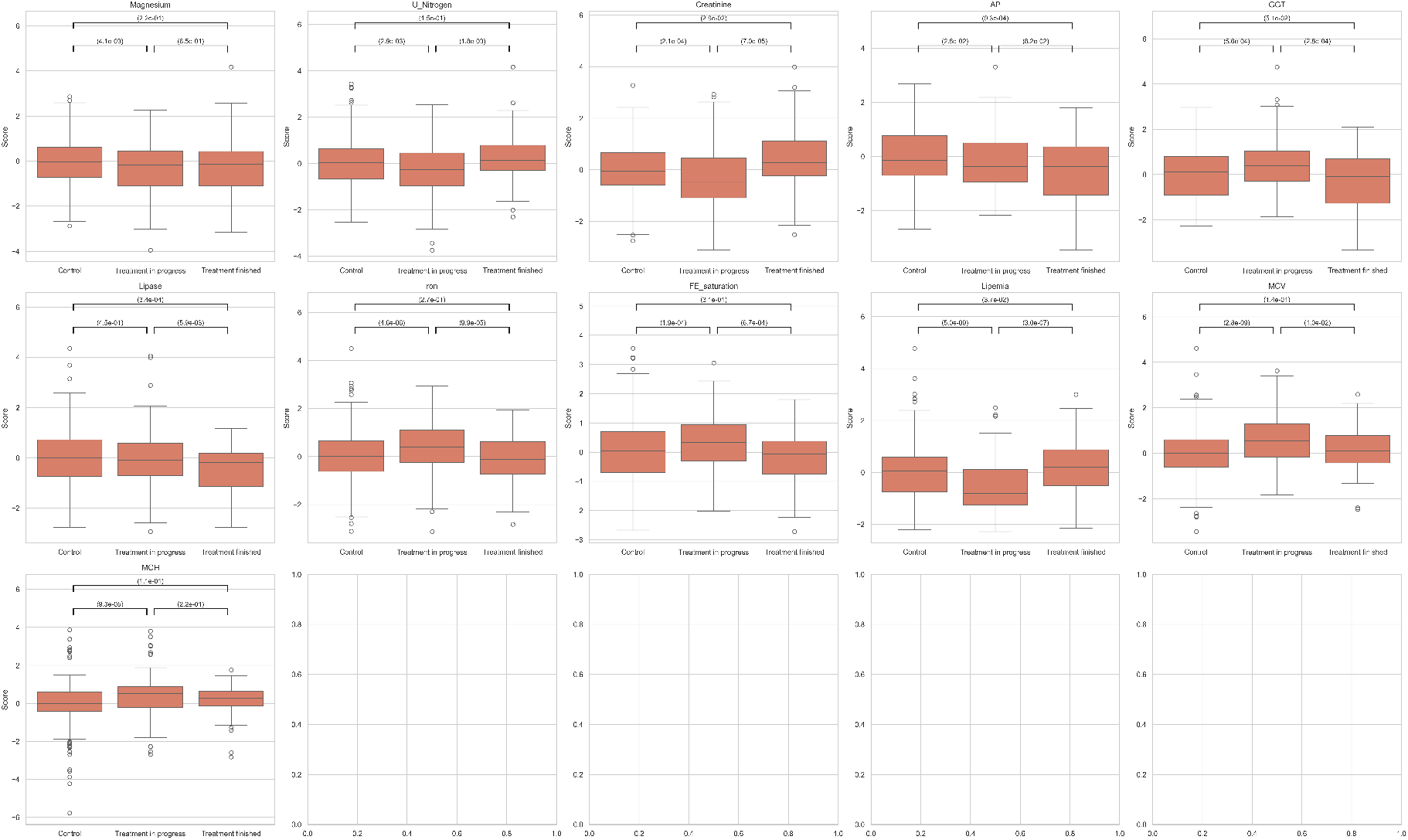
Individual blood markers changed in response to LAM (only *p/*15 *<* 0.05 are shown) in groups of the animals before, during and after the treatment.

**FIG. S3:**
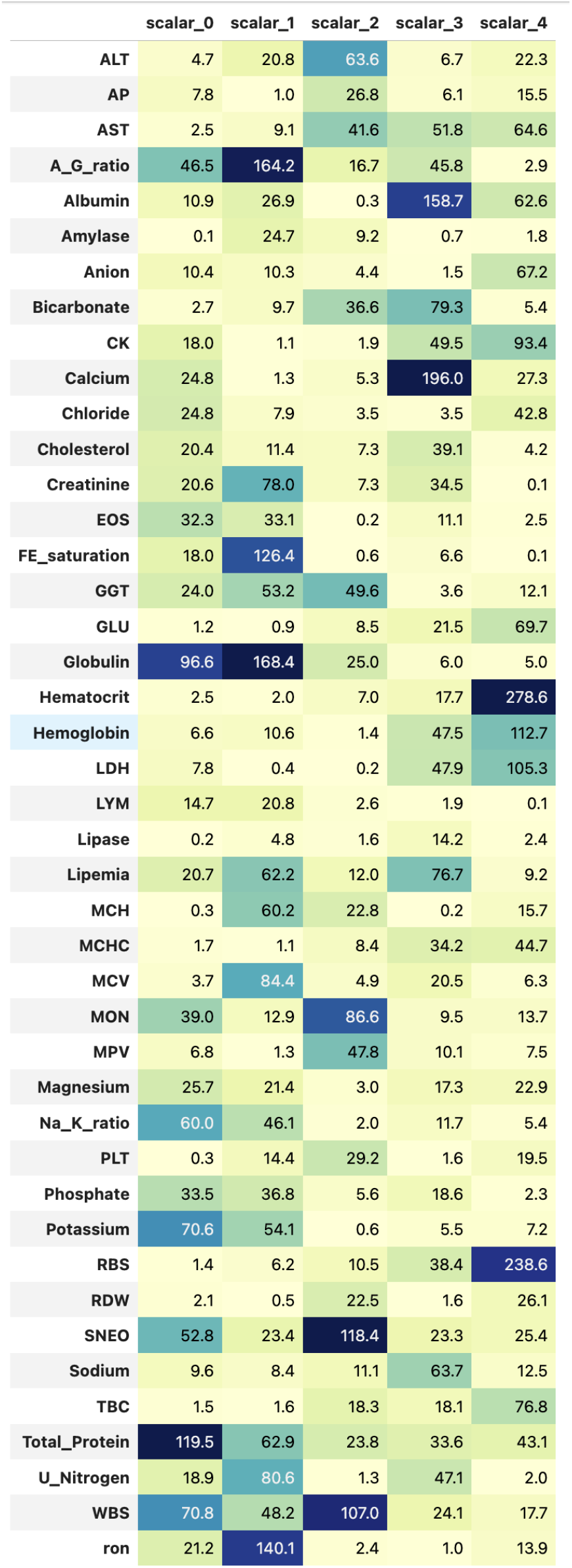
Association of LHM’s latent variables with blood biomarkers

**FIG. S4:**
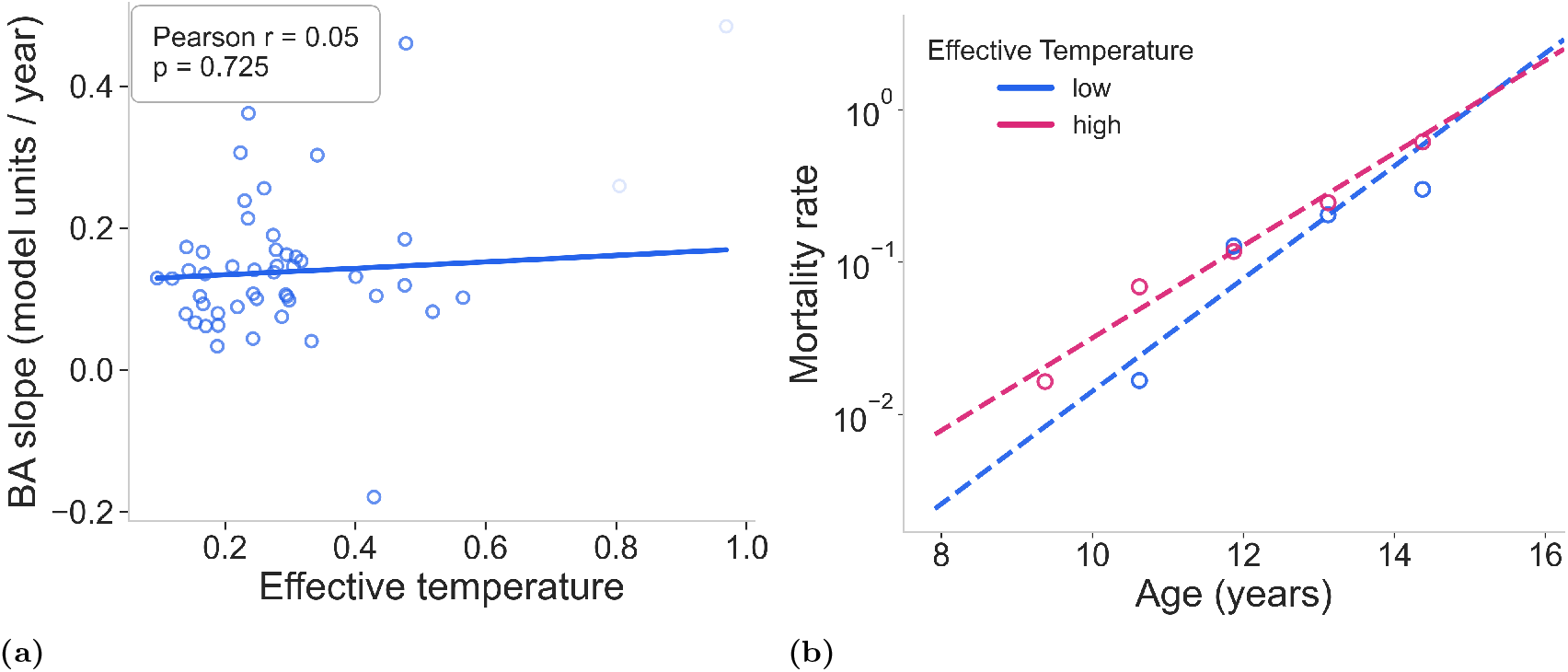
(a) Damage accumulation rate, quantified by the slope of model-derived Biological Age (BA) trajectories, as a function of the effective phenotypic temperature. (b) Empirical mortality hazard rates calculated in 15-month bins for canine cohorts stratified by median phenotypic temperature (Cold, blue; Hot, red). Points represent empirical hazard rates (deaths per animal-month), and solid lines indicate the Gompertzian fits (straight lines in a log-scale) for ages *>* 60 months.

**FIG. S5:**
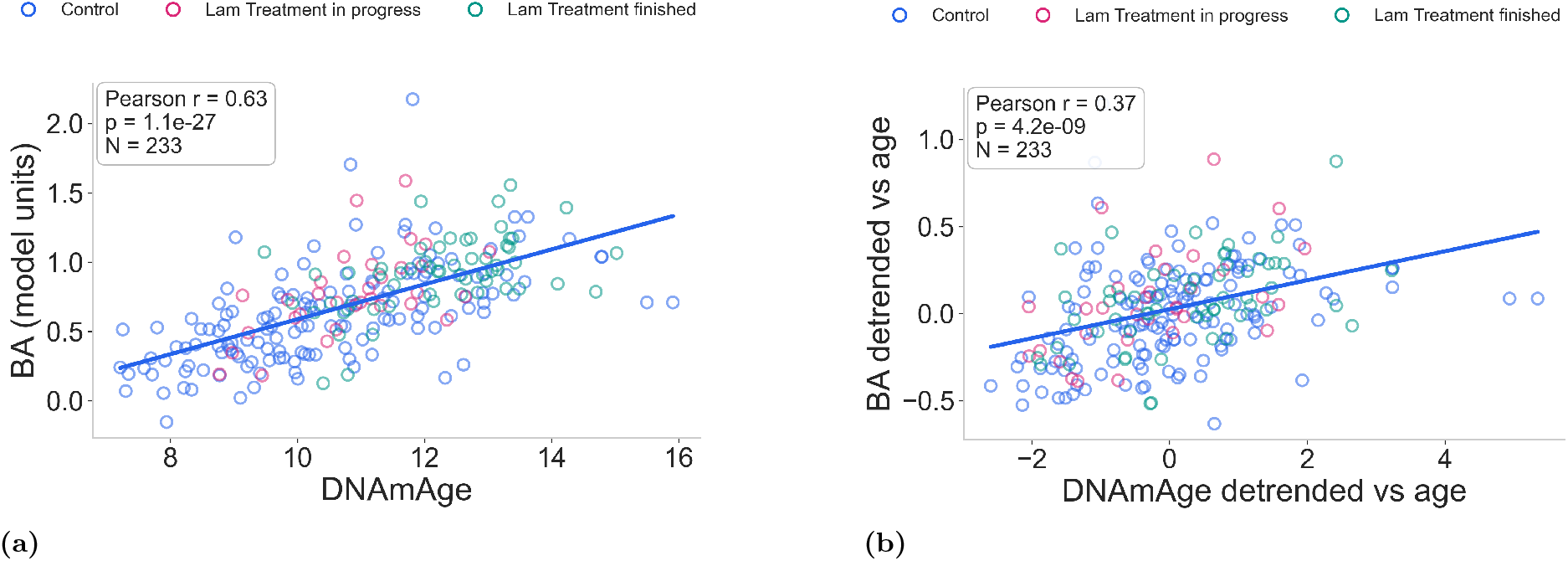
Biological age from DNAmAge vs. the BA from LHM **(A)**; biological age from DNAmAge vs. the BA from LHM where both are detrended from the common dependence on chronological age **(B)**.

**FIG. S6:**
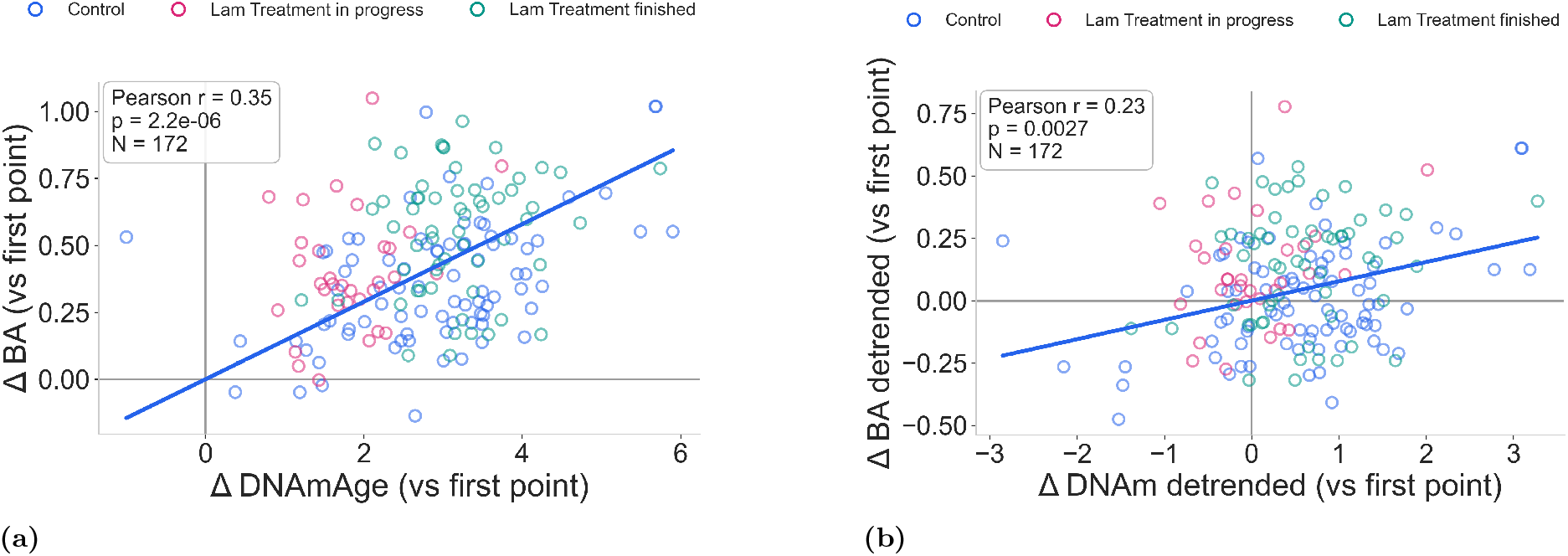
Biological age from ∆DNAmAge vs. the ∆BA from LHM (∆ is vs. the first available point per dog) **(A)**; biological age from ∆DNAmAge vs. the ∆BA from LHM where both are detrended from the common dependence on chronological age **(B)**.

